# EraSOR: Erase Sample Overlap in polygenic score analyses

**DOI:** 10.1101/2021.12.10.472164

**Authors:** Shing Wan Choi, Timothy Shin Heng Mak, Clive J. Hoggart, Paul F. O’Reilly

## Abstract

**Background:** Polygenic risk score (PRS) analyses are now routinely applied in biomedical research, with great hope that they will aid in our understanding of disease aetiology and contribute to personalized medicine. The continued growth of multi-cohort genome-wide association studies (GWASs) and large-scale biobank projects has provided researchers with a wealth of GWAS summary statistics and individual-level data suitable for performing PRS analyses. However, as the size of these studies increase, the risk of inter-cohort sample overlap and close relatedness increases. Ideally sample overlap would be identified and removed directly, but this is typically not possible due to privacy laws or consent agreements. This sample overlap, whether known or not, is a major problem in PRS analyses because it can lead to inflation of type 1 error and, thus, erroneous conclusions in published work.

**Results:** Here, for the first time, we report the scale of the sample overlap problem for PRS analyses by generating known sample overlap across sub-samples of the UK Biobank data, which we then use to produce GWAS and target data to mimic the effects of inter-cohort sample overlap. We demonstrate that inter-cohort overlap results in a significant and often substantial inflation in the observed PRS-trait association, coefficient of determination (R^2^) and false-positive rate. This inflation can be high even when the absolute number of overlapping individuals is small if this makes up a notable fraction of the target sample. We develop and introduce EraSOR (<u>Era</u>se <u>S</u>ample <u>O</u>verlap and <u>R</u>elatedness), a software for adjusting inflation in PRS prediction and association statistics in the presence of sample overlap or close relatedness between the GWAS and target samples. A key component of the EraSOR approach is inference of the degree of sample overlap from the intercept of a bivariate LD score regression applied to the GWAS and target data, making it powered in settings where both have sample sizes over 1,000 individuals. Through extensive benchmarking using UK Biobank and HapGen2 simulated genotype-phenotype data, we demonstrate that PRSs calculated using EraSOR-adjusted GWAS summary statistics are robust to inter-cohort overlap in a wide range of realistic scenarios and are even robust to high levels of residual genetic and environmental stratification.

**Conclusion:** The results of all PRS analyses for which sample overlap cannot be definitively ruled out should be considered with caution given high type 1 error observed in the presence of even low overlap between base and target cohorts. Given the strong performance of EraSOR in eliminating inflation caused by sample overlap in PRS studies with large (>5k) target samples, we recommend that EraSOR be used in all future such PRS studies to mitigate the potential effects of inter-cohort overlap and close relatedness.

## Introduction

Polygenic risk scores (PRSs) are proxies of individuals’ genetic liability to a trait or disease [1] that have been applied in numerous research settings, including patient stratification [2] and investigation of treatment response [3–6]. The power of PRS analyses is dependent on the heritability and polygenicity of the trait, the power of the genome wide association study (GWAS) used to derive the PRS, and the size of the target data sample [7]. The recent surge of high quality genetic and phenotypic data from large-scale biobank projects, such as the UK Biobank [8], BioBank Japan [9], Taiwan Biobank [10], and FinnGen [11], as well as GWAS resources from large consortia such as the Psychiatric Genomic Consortium (PGC) [12], GIANT [13] and the Global Lipids Genetics Consortium (GLGC) [14] have provided unprecedented opportunity to perform highly-powered PRS analyses.

However, expansion in data size does not come without a cost in this setting: as sample sizes increase, there is greater risk that samples are recruited into multiple cohorts or that entire cohorts are included in multiple consortia. For PRS analyses, which typically test for association between PRS and a trait(s) or outcome of interest, overlapping samples between the GWAS and target data samples can result in spurious inflation of the coefficient of determination (*R*^*2*^) and association *P*-values, leading to false-positive and exaggerated findings [15]. Overlapping samples should ideally be removed from either the GWAS or target data to avoid misinterpretation of results, but participant privacy agreements usually limit access to raw genotyping data, meaning that this is generally not an option.

Here we first evaluate the extent to which different degrees of sample overlap and relatedness between GWAS and target samples generates biased PRS-trait associations. Next, to overcome the sample overlap problem, we develop and introduce EraSOR (Erase Sample Overlap and Relatedness), a python software that adjusts GWAS summary statistics [1] to correct for inflation of PRS-trait association results caused by overlapping samples between the GWAS and target samples. Through extensive simulations using the UK Biobank genetic data [8], we demonstrate that EraSOR can robustly adjust for inflation in test statistics caused by various degrees of overlapping samples, level of relatedness, or ascertainment schemes in case/control settings. We propose that EraSOR will increase the accuracy of results in all future PRS studies with known sample overlap and will act as a sensitivity tool for assessing the reliability of results in PRS studies with unknown but potential sample overlap. EraSOR is an open-source software and is freely available at **https://gitlab.com/choishingwan/EraSOR**.

## Methods

### EraSOR framework

Consider two GWAS *k* = {1, 2} performed on the same continuous outcome *Y*_*k*_. The effect size of the *g*^th^ SNP in study *k* (*β*_*kg*_) is estimated using a regression model

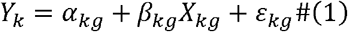

where *X*_*kg*_ is the standardized genotype vector for SNP *g* in study *k*, and *ε*_*kg*_ is the random error assumed to be independent between studies. Under the null model of no contribution of SNP *g* to the trait, *β*_*kg*_ *=* 0, and assuming no sample overlap, then 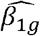 and 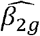 estimated from the two GWASs should be independent, i.e.,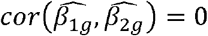. However, when there are overlapping samples between the two studies, then a correlation is induced between the regression coefficients, such that 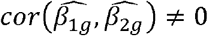. From LeBlanc et al [16], this correlation can be approximated as

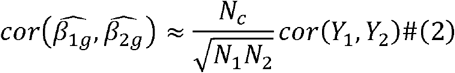

for quantitative traits, where *cor* (*Y*_1_,*Y*_2_) represents the correlation between the traits; *N*_*c*_ is the number of overlapping samples; and *N*_1_, *N*_2_ are the sample sizes of studies 1 and 2, respectively [16]. Since we are considering only a single phenotype here, *cor* (*Y*_1_,*Y*_2_) is equal to 1, and so we have:

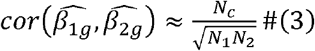

which captures correlations only due to sample overlap independent of the true causal effect. Assuming sample overlap does not affects the standard error estimates, LeBlanc et al [16] proposed that when the number of overlapping samples (*N*_*c*_) is known, one can adjust the joint distribution of the summary statistics (z-scores) of the two GWASs as:

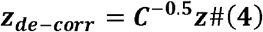

where ***z*** is a 2-by-*M* matrix containing z-scores estimated in each study, *M* is the number of SNPs common to both studies, and ***C*** is the 2×2 matrix with ones on its diagonal and 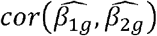 as its off-diagonal elements. While this adjustment is effective [16], it requires prior knowledge of *N*_*c*,_ which is typically unknown in PRS studies. However, we propose utilizing univariate and bivariate LD score regression [17,18] to estimate 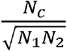 and thus 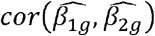 from Eq. 3 as follows:

Bivariate LD score regression is typically used to estimate the genetic correlation between two traits using the GWAS corresponding to each, and is defined in [13] by the following equation:

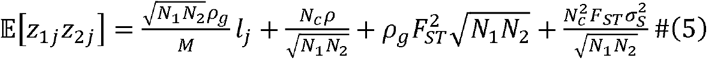

where *l*_*j*_ is the LD score of SNP *j*; *ρ*_*g*_ is the genetic covariance between the two traits; *ρ* = *ρ*_*g*_ + *ρ*_*e*_ ; *ρ*_*e*_ is the non-genetic covariance; *F*_*ST*_ and *σ*_*s*_ are the genetic and environmental stratification respectively.

LDSC assumes two underlying populations within each cohort, and that the levels of genetic and environmental stratification are similar in the two cohorts, e.g. 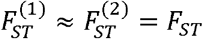 and *σ*_*S*1_ ≈ *σ*_*S*2_ = *σ*_*S*_ [13]. Since we are considering only a single phenotype here, *ρ* is equal to 1, and so we have:

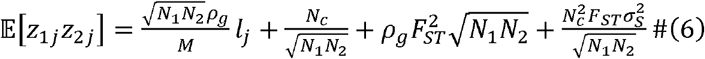

We wish to solve for N_c_ and hence apply Eq. 3 to generate a de-correlated base GWAS that does not lead to inflated PRS-trait associations due to sample overlap. To do this, we will utilize the univariate LD score regression model. The univariate LD score regression equation can be derived as a special case of the bivariate LD score equation by assuming that the two outcomes and cohorts are identical [13,17], leading to:

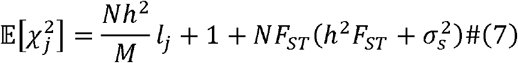

Univariate LD score regression performs a regression of observed *χ*^2^ on *l*_*j*_, with the effect size estimate of *1*_*j*_ corresponding to a scaled estimate of heritability 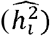 and with the estimated intercept term, 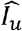 as follows:

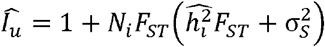

If we assume that the environmental stratification 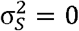, then we have:

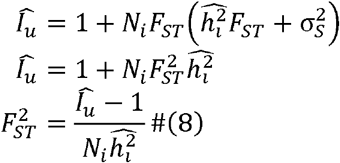

Since we can estimate 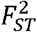 using both the base and target data, we then take the weighted mean estimate of both:

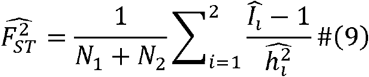

The intercept term of the bivariate LD score regression is:

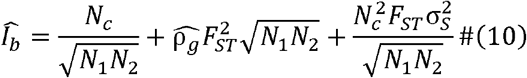

Substituting Eq. 9 and 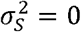 into Eq. 10, we have:

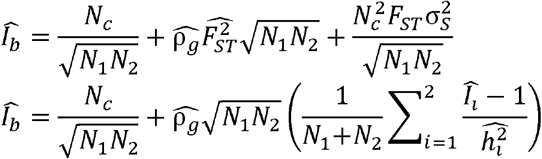

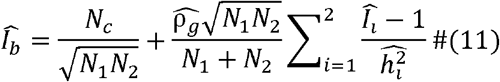

Since we can estimate the genetic covariate 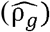, the trait heritability 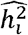 and the intercepts from the univariate and bivariate LD score regression analyses of the GWAS and target data, we can obtain an estimate of 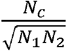. Substituting this estimate into Eq. 3 will derive an estimate of 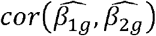 that can be used to produce de-correlated GWAS z-statistics via Eq.4. EraSOR automatically performs the bivariate LD score and univariate LD score regression analyses on the GWAS summary statistics generated from the base and target data. To test the performance of EraSOR, including its robustness to the modelling assumptions (e.g., assuming 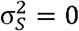), we performed a series of extensive simulations.

### UK Biobank genotype data

The UK Biobank is a prospective cohort study of around 500,000 individuals recruited across the United Kingdom during 2006-2010. The genetic data from UK Biobank comprises 488,377 samples and 805,426 SNPs. Standard quality control (QC) procedures were performed, removing any SNPs with minor allele frequency < 0.01, genotype missingness > 0.02 and with a Hardy Weinberg Equilibrium Test *P*-value < 1×10^−8^. Samples with high levels of missingness or heterozygosity, with mismatching genetic-inferred and self-reported sex, or with aneuploidy of the sex chromosomes were removed as recommended by the UK Biobank data processing team. Next, 4-means clustering was applied to the first two Principal Components (PCs) of the genotype data and those individuals in the (largest) cluster corresponding to European ancestry were retained for the primary analyses because polygenic risk scores have been shown to have low portability between ancestries [14] motivating ancestry-matched PRS studies until cross-ancestry PRS methods are developed, which our main results correspond to (see section *Samples with population stratification* below, which describes analyses that we also performed on individuals of all ancestries in the UK Biobank). A greedy algorithm [19] was then used to remove related individuals, with kinship coefficient > 0.044, in a way that maximized sample retention. In our simulations that investigate the effect of related individuals in the GWAS and target data, we instead randomly retain one first degree relative (defined as kinship coefficient ≥ 0.177 and ≤ 0.354) of a randomly sampled individual in the GWAS data. Altogether, we retain 557,369 SNPs, 387,392 individuals and 23,429 of their first-degree relatives for the set of analyses performed. For the simulations of population stratified samples, we extracted samples 10 standard deviations from the centroid of the European cluster and defined these as “non-European” samples. Quality control procedures were repeated using the parameters described above after combining these non-European samples with the European samples, resulting in 387,365 samples of European ancestry and 21,779 individuals of non-European ancestry. Code used to perform the QC and corresponding documentation are available at **https://choishingwan.gitlab.io/ukb-administration/admin/master_generation/**. This research has been conducted using the UK Biobank Resource under application 18177 (Dr O’Reilly).

### Phenotype simulation

#### Quantitative Traits without population structure

Quantitative phenotypes (*Y*) with heritability (*h*^*2*^) of 0, 0.1, and 0.5 were simulated using the UK Biobank genotype data (post QC; see above) as input. Quantitative traits were simulated as:

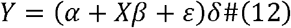

where *X* is the standardized genotype matrix corresponding to all samples and 10,000 randomly selected SNPs with effect size *β* following a standard normal distribution. *Xβ* were adjusted such that it has mean 0 and variance of *h*^*2*^; and *ε* represents the random error, which follows 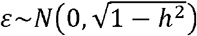. To ensure EraSOR works for distribution that are not only standard normal, we included α as the phenotypic mean randomly sampled from a normal distribution with mean 0 and standard deviation of 1, and δ as the phenotypic variable randomly sampled from 1 to 100 to simulate phenotypes that does not follow the standard normal distribution.

To model polygenic risk score analyses with sample overlap, we randomly selected either 120k or 250k individuals from the sample of 387,392 individuals available to us (see above) to generate two different sizes of base GWAS data. Next, we randomly sampled 1,000, 5,000 or 10,000 individuals from the remaining sample to act as three different sizes of target data, of which 0%, 5%, 10%, 50% or 100% were randomly selected from the base data sample so that there was a known degree of sample overlap between the base and target data. In addition, we generated an “overlap-free” base cohort in which the overlapping samples were removed from the base cohort so that we could compare the result of applying EraSOR against results of physically removing overlapped samples from the base cohort.

In order to search a feasible parameter space in sufficient depth, we only simulate phenotype with heritability of 0.5, with a base cohort of 250k and target cohort of 5,000; only simulate base cohort with 120k samples when the phenotypic heritability is ≤ 0.1 and target cohort has 5,000 samples; and only simulate target cohort with 1,000 and 10k samples when the base cohort contain 250k samples and the phenotypic heritability is ≤ 0.1. The entire set of simulations were repeated 100 times.

#### Binary Trait

Binary traits were simulated under the liability threshold model [20], simulating a normally distributed liability using Eq. 12 with α = 0, δ = 1, and cases defined as samples with disease liability higher than liability thresholds of 0.9, 0.7 and 0.5, corresponding to population prevalences of 0.1, 0.3 and 0.5, respectively. To limit the complexity of our simulations, the sample prevalence of our cohorts follows the population prevalence.

In the binary trait setting, overlap can be ascertained such that the overlap is among cases, or among controls, or among both. To investigate the effect of case-only or control-only overlap, we randomly selected 120k effective samples (effective samples defined as *N*_*eff*_ = 4/(1/*N*_*cases*_ + 1/*N*_*controls*_) [21]) as the base cohort, and then randomly selected 5,000 effective samples as the target cohort, where 0%, 5%, 10%, 30% or 50% of the cases or of the controls in the target cohort were sampled from the base cohort. We also performed simulations where the overlapping samples were selected at random among cases and controls. An “overlap-free” base cohort was generated with all overlapping samples removed.

In order to search a feasible parameter space in sufficient depth, we only vary the trait heritability when the population prevalence is 0.1, and only vary the population prevalence when the trait heritability is ≤ 0.1. These simulations were repeated 100 times.

### Related samples

Spurious inflation in PRS analysis test statistics may also be observed when there are closely related individuals between the base and target cohorts. To investigate the effects of relatedness on PRS results, we repeated the quantitative trait simulations with a modified Eq. 12:

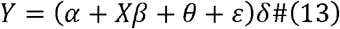

where *θ* is the shared environment between the related individuals and follows a random normal distribution with mean 0 and variance 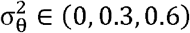 if and only if 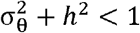, with each related pair of individuals having the same *θ* value. *ε* represents a combination of non-shared environment and random error, which follows 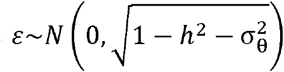. To model the inter-cohort relatedness, we first select all individuals with a first-degree relative in the UK Biobank (kinship coefficient ≥ 0.177 and ≤ 0.354), of which there are 23,429 individuals, and then randomly select additional samples who do not have any first-degree relatives to form a base cohort containing 250k samples. We then generate target cohorts containing 5,000 samples, with either 0%, 30%, 60% or 100% of the target samples being first-degree relatives of samples in the base cohort. We also generated a reference cohort from the base cohort where all the related samples in the target cohort were replaced by unrelated individuals for benchmarking the performance of EraSOR. The entire set of simulations were repeated 100 times.

### Samples with population stratification

An assumption of the EraSOR algorithm is that the environmental stratification 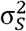 is zero. When environmental stratification is present, 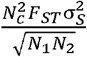 from Eq. 10 is no longer 0 and a bias proportional to the environmental stratification and the genetic stratification (*F*_*ST*_) may be introduced. We devised two strategies for simulating data with both environmental and genetic stratification to test the sensitivity of EraSOR to deviations of each from 0. In the first, we partitioned the UK Biobank into European and non-European ancestries, while in the second we used the simulation software HapGen2 [22].

In the first simulation strategy, the UK Biobank samples were divided into European and non-European ancestries based on 4-mean clustering on PC1 and PC2 (see above). Quantitative traits with environmental stratification were then simulated as:

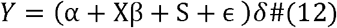

with the environmental stratification term (S) defined as

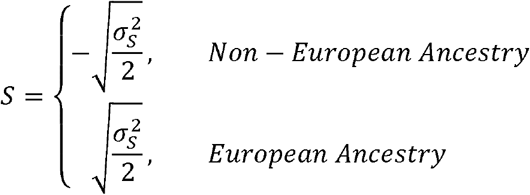

where 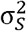 can take a value of 0, 0.3 or 0.9 if and only if 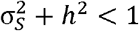, and *ϵ* represents the residual term, which follows 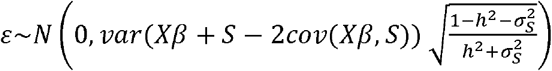, with cov(Xβ,S) being the covariance between Xβ and S. To investigate the effect of sample overlap in the presence of environmental and genetic stratification, we randomly selected either 120k or 250k individuals from the sample of 409,144 individuals available to us (see above) to generate two different sizes of base GWAS data. Next, we randomly sampled 5,000 or 10,000 individuals from the remaining sample to act as two different sizes of target data, of which 0%, 10%, 50% or 100% were randomly selected from the base data sample. To ensure that the genetic and environmental stratification is the same within the base and target data, the same ancestry ratio was maintained in all simulated data sets, matching the ratio in the full data set (∼5% non-European ancestry). In addition, we generated an “overlap-free” base cohort in which the overlapping samples were remove from the base cohort to allow benchmarking the performance of EraSOR. The entire set of simulations were repeated 25 times.

Given that only ∼5% of the UK biobank samples correspond to individuals of non-European ancestry, the effect of genetics and environmental stratification may be limited. Thus, we developed a second strategy to test their effects in which we used HapGen2 [22] to simulate 180k Yoruban and 180k Finnish samples using recombination maps from the 1000 Genomes Project [23]. 500 “Finnish” samples and 500 “Yoruban” samples were selected to calculate the LD scores using LDSC (v1.0.1) and flashPCA (v2.0) [24] was used to calculate the first 15 PCs of the data.

We repeated the population stratification simulation using the HapGen2 simulated genotype data, with S represented now segregate according to the simulated population. The entire set of simulations were repeated 25 times.

### Genome Wide Association Study and Polygenic Score Analysis

Genome wide association analyses (GWAS) were performed on the base and target cohorts using PLINK 2.0 (version 2021-08-04) [25] with the *--glm* function. As binary traits were only simulated for the European ancestry only analyses, where population structure was not simulated, and considering the computational cost of including covariates in the logistic regression, we did not include PCs in our binary trait analysis. On the other hand, quantitative traits were simulated in all scenarios, some of which are population stratified. Thus, we included 15 PCs as a covariate for our quantitative triat analyses. The resulting summary statistics were then provided to EraSOR to generate the adjusted summary statistics using European LD scores [17] calculated from 1,000 Genomes Project Phase 3 data [23] or the LD scores calculated from a subset of the simulated genotypes (HapGen2 simulation) using LDSC (v1.0.1) [17]. PRS analyses using the adjusted, unadjusted, and the “overlap-free” summary statistics were performed using PRSice-2 (v2.3.5) [26] with the default settings. The R^2^ and *P*-value of association of the PRS-trait tests were reported.

### Strategy for Benchmarking

To investigate the level of spurious inflation caused by inter-cohort relatedness and overlapped samples, we first established a baseline PRS R^2^, calculated using base cohorts without overlapped samples. The bias can then be measured as the observed PRS R^2^ minuses the baseline PRS R^2^ (Δ*R*^2^), given the same phenotype and cohort sizes. For non-heritable traits, we also measure the level of false-positive, defined as any PRS with *P*-value < 1×10^−4^ [27].

On the other hand, to compare the performance of EraSOR with the optimal strategy of directly removing overlapping samples – an option that is typically not available – we calculate PRSs: (i) using summary statistics adjusted by EraSOR (“adjusted PRS”) and (ii) using summary statistics generated from a base cohort with all overlapping and/or related samples removed (“overlap-free PRS”). We present the performance of EraSOR as the PRS-trait association R^2^ of the adjusted PRS minuses the R^2^ of the overlap-free PRS R^2^ (Δ*R*^2^). If EraSOR has successfully corrected for the sample overlap, then Δ*R*^2^ should be close to 0.

## Results

### Inflation caused by overlap

The presence of overlapping samples between the base and target data sets is known to cause inflated association between polygenic risk scores (PRS) and phenotypes [15], but the extent and characteristics of the problem have not been described. Here, we performed extensive simulations using the UK Biobank [8] genotype data to investigate the inflation caused by different levels and types of inter-cohort sample overlap in relation to traits simulated with varying heritability and prevalence (see Methods). Base and target cohorts were generated with varying degrees of sample overlap, measured as 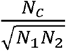, where *N*_*c*_ is the number of overlapping samples and *N*_1_ and *N*_2_ are the sample sizes of the base and target cohort, respectively. PRS analyses were conducted using the standard *clumping+thresholding* (C+T) PRS calculation method [1], implemented in *PRSice* [26].

We first estimated the false-positive rate induced by sample overlap by simulating non-heritable traits and recording the fraction of significant PRS-trait association (Supplementary Fig. 1). Highly significant associations between PRS and non-heritable phenotypes were observed when even limited inter-cohort sample overlap was present (Fig. 1). Specifically, for non-heritable quantitative traits, the inflation in association (e.g. *p-*value of association) is highly positively correlated with the degree of overlap (Pearson Correlation coefficient (*γ*) = 0.96, *P*-value < 2.2×10^−16^). For example, when there is a base cohort of 250k samples, target cohort of 5,000 samples and 250 overlapping samples (5% of target sample; degree of overlap = 0.0071) the false positive rate is 17%, while this increases to 94% when there are 500 overlapping samples (10% of target sample; degree of overlap = 0.014) (Fig 1a).

**Figure 1.**
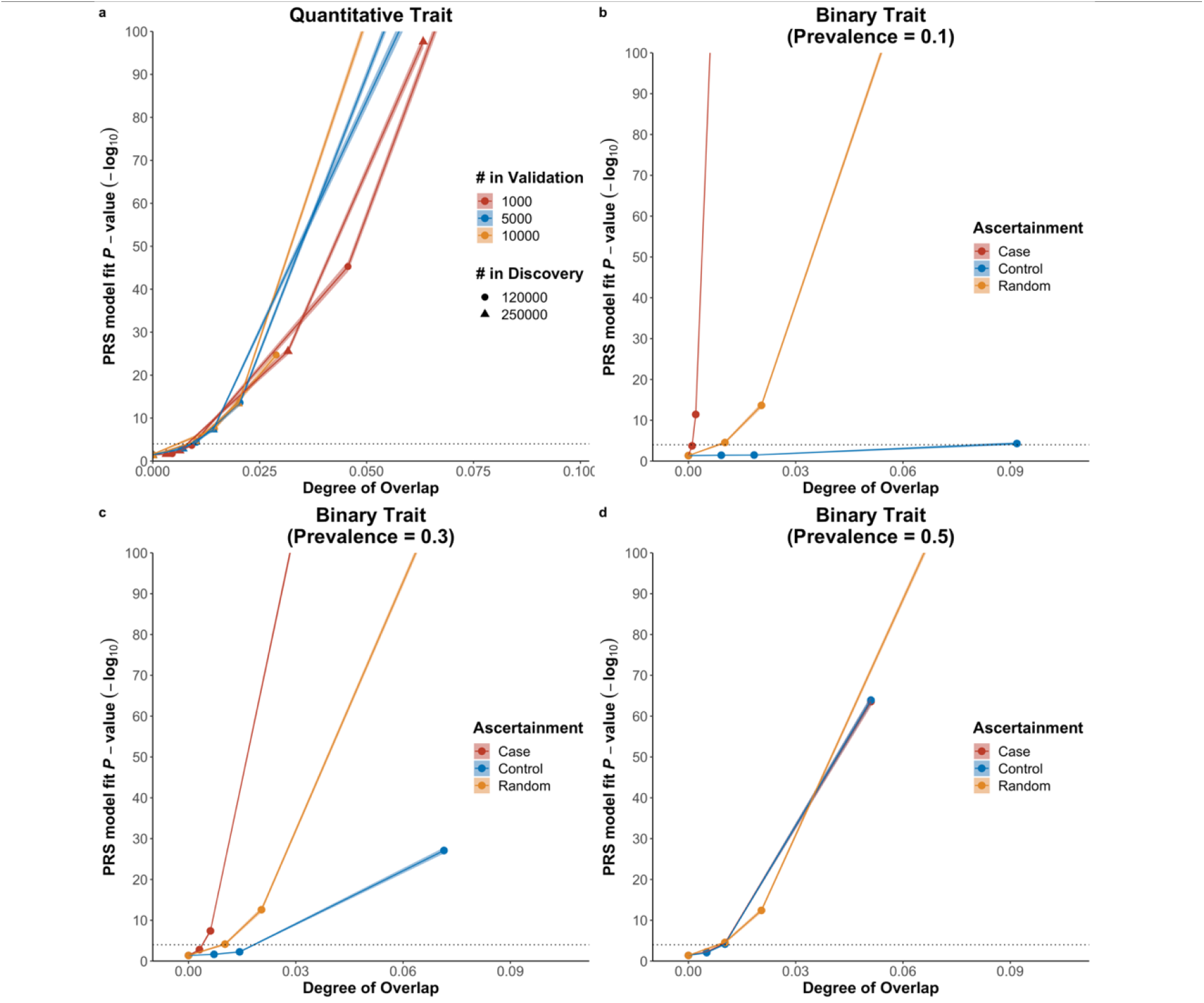
Effect of sample overlap on performance of PRS for non-heritable traits. The dotted line represents the significance threshold *(P-*value = 1×10^−4^) for high-resolution testing in *PRSice* [27]. The X-axis shows the degree of overlap, calculated as 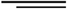 and the Y-axis shows the -log_10_ transformed p-value of association between the PRS and the simulated phenotype. Shaded area represents the 95% confidence interval. However, as the confidence interval is small, it is difficult to observe on the scale of these plots **a)** Quantitative traits with different cohort sizes. **b)** Binary traits with population prevalence of 0.1 **c)** Binary traits with population prevalence of 0.3 **d)** Binary traits with population prevalence of 0.5

In the binary trait setting, sample overlap may be among cases only, controls only, or be among both. These alternatives were investigated by first simulating binary traits with different population prevalence using the liability threshold model [20]. Cohorts with effective sample sizes of 120k in the base data and 5000 in the target data were generated with different degrees and scenarios of sample overlap. We observed extreme inflation associated with case-only overlap when population prevalence is lower than 0.5. For a binary trait with population prevalence 0.1, a false positive rate of 43% is observed when the degree of overlap is 0.001, which corresponds to 5% of the cases from the target cohort also present in the base cohort (Fig 1b). When the degree of sample overlap is doubled (∼0.002), the false positive rate is 100%. The inflation in PRS-trait association is not as sensitive to control-only sample overlap when the population prevalence is small. We observe a false positive rate of ∼47% when the degree of overlap is as high as 0.092, which corresponds to 50% of the controls from the target cohort also present in the base cohort (Fig 1b). This discrepancy between the effect of case and control overlap is a result of the differential contribution of cases and controls to the PRS-trait association in our simulations. Cases are sampled from the extreme upper tail of the liability distribution at a frequency corresponding to the disease prevalence, which is typically low: this gives each case greater weight in the calculation of the PRS-trait association and, thus, an overlapping case will generate greater inflation than an overlapping control. This was consistent with our simulation results (Fig 1c, 1d), where the inflation in Δ*R*^2^ caused by overlapping cases decreases as population prevalence increases (*γ*= −0.068, *P*-value = 4.70×10^−5^). The reverse relationship between inflation and population prevalence was observed for control-only overlap (*γ* = 0.20, p-value = 3.90×10^−34^). For a population prevalence of 0.5, case-only and control-only overlap have the same impact on the inflation (Fig 1d).

Given that complete overlap of individuals in the base and target data can generate PRS-trait associations that are severely inflated, closely related individuals independently enrolled into the base and target cohorts may induce some inflation considering their shared genetics and environment. Here we tested the effect of relatedness between the base and target cohorts on PRS-trait associations in non-heritable traits in a similar way to that for sample overlap (see Methods), where inter-cohort relatedness is defined as 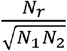 where *N*_*r*_ is the number of samples in the target cohort that are first degree relatives with samples in the base cohort. A false positive rate of 76% is observed when the inter-cohort relatedness is 0.085 and the shared environment explains 30% of the trait variance (250k base, 5k target, 60% of target samples have 1^st^ degree relatives in the base), while a false positive rate of 84% is observed with an inter-cohort relatedness of 0.042 for traits with shared environment contribution of 60% (30% related samples). See Supplementary Fig. 2 for full results regarding the effects of inter-cohort relatedness on PRS-trait association inflation.

In the next section we extend these investigations to consider the effects of sample overlap on PRS-trait associations on heritable traits, but we present these findings in conjunction with results based on the application of our method EraSOR, which is designed to resolve the problem.

### Performance of EraSOR

To tackle the problem of inflation caused by inter-cohort overlap and relatedness, we developed the <u>Era</u>se <u>S</u>ample <u>O</u>verlap and <u>R</u>elatedness (EraSOR) method. Using GWAS summary statistics generated from the base and target cohorts, EraSOR implements univariate and bivariate LD score regression [17,18] to estimate several parameters that are then used to perform a de-correlation calculation of the base GWAS test statistics (see Methods). These adjusted base GWAS summary statistics can then be used for downstream PRS analyses, with sample overlap or relatedness corrected for.

In order to evaluate the performance of EraSOR, we conducted an extensive set of simulations covering a range of scenarios of inter-cohort sample overlap and relatedness (see below and Methods).

### Simulations using UK Biobank data

We observed that for both quantitative and binary traits, EraSOR almost entirely eliminates the inflation caused by inter-cohort overlap and relatedness in our simulations based on UK Biobank (European ancestry base and target samples) data (Figure 2). These simulations modelled a range of scenarios that varied trait heritability, prevalence, degree of overlap and combinations of overlap among cases and controls. For example, simulating quantitative traits with heritability 0.1, a base cohort of 250k samples, target cohort of 5000 samples, and degree of overlap 0.141 – in which all samples in the target data are also in the base GWAS – the mean Δ*R*^2^ is -3.76×10^−4^ (standard error: 3.03×10^−4^). Left unadjusted, the mean Δ*R*^2^ is approximately 0.35 (standard error 0.00125), suggesting that EraSOR has removed the inflation introduced by sample overlap. A similar pattern of complete removal of the effects of sample overlap is observed for the quantitative traits across the full range of heritability and cohort sample sizes tested (Figure 2a).

**Figure 2.**
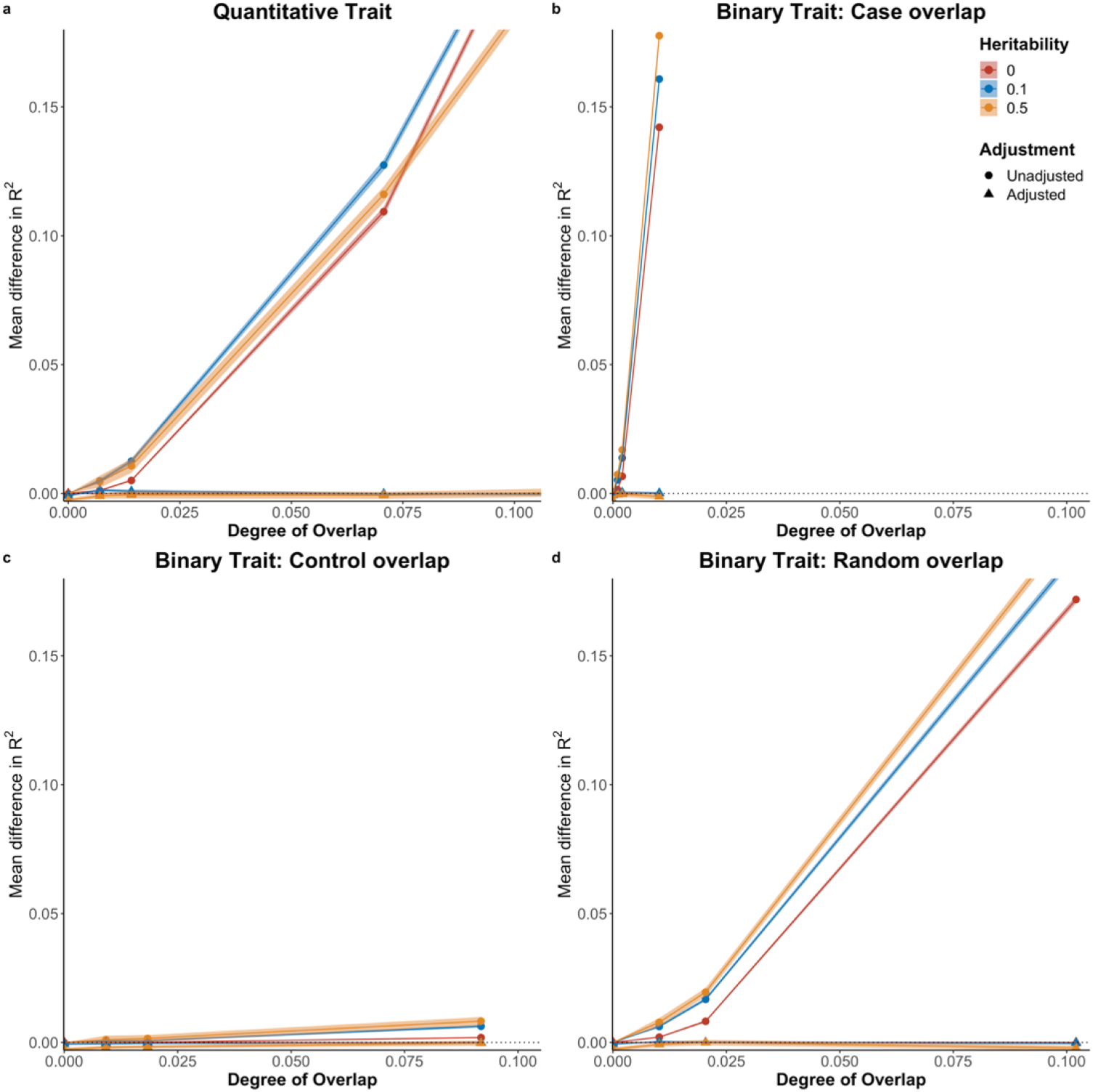
Comparing the performance of the PRS using the EraSOR adjusted summary statistics and the unadjusted summary statistics. The X-axis shows the degree of overlap, and the Y-axis shows the mean difference between the observed R^2^ and the expected R^2^. Shaded area represents the 95% confidence interval (small on this scale). **a)** Performance in quantitative traits with 250,000 samples in the base cohort and 5,000 samples in the target cohort; b) Performance in binary traits with prevalence of 0.1 and where overlap samples were ascertained for cases; **c)** ascertained for controls; **d)** or were randomly ascertained.

**Figure 3.**
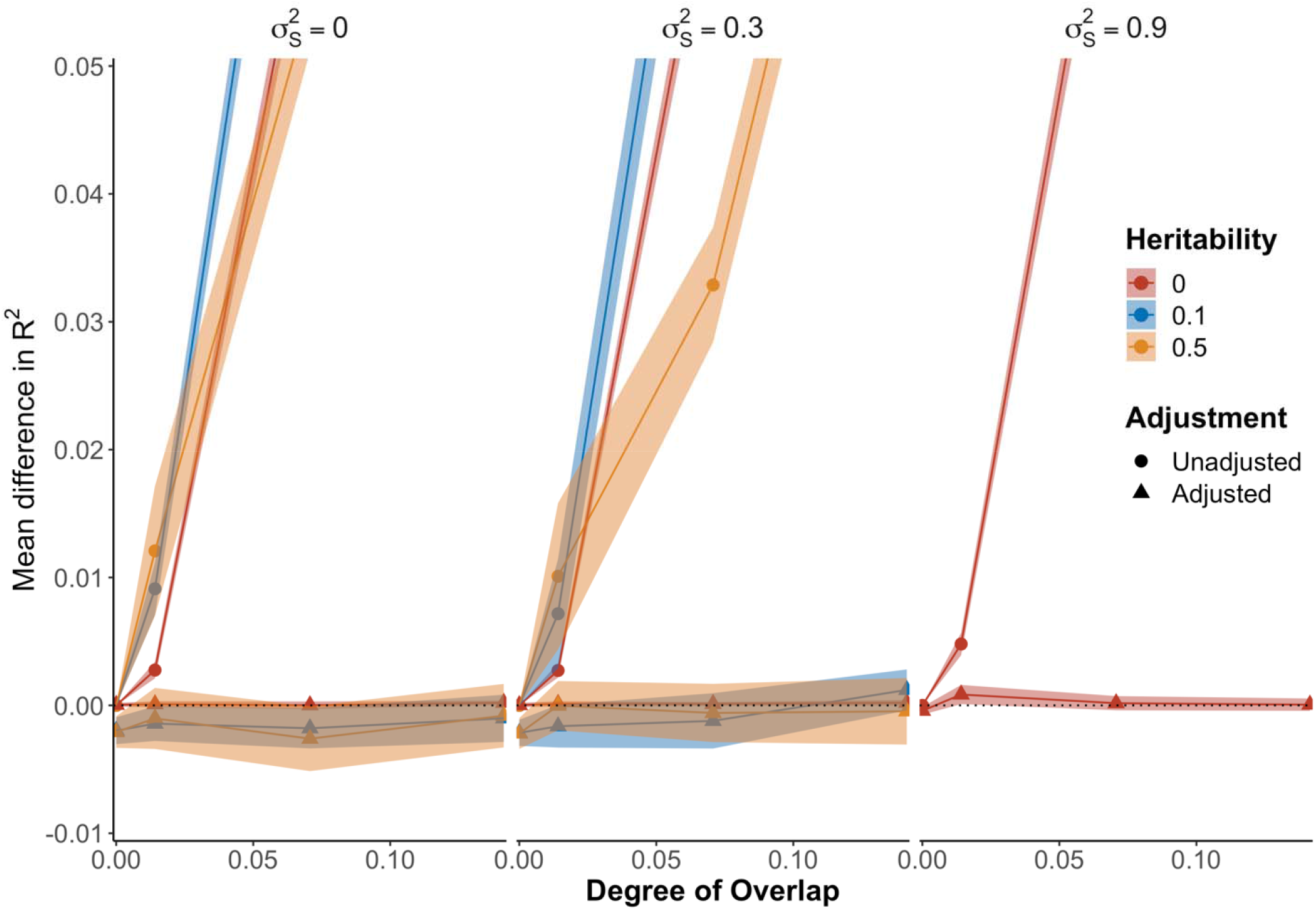
Performance of EraSOR from the HapGen2 simulation. Y-axis represents the mean difference between the observed R^2^ and the expected R^2^ and X-axis represents the degree of overlap. Range shows the 95% confidence interval. Based on simulation results, it seems like EraSOR are robust against different environmental stratification ().

EraSOR also performs extremely well for binary traits (Figure 2b, 2c, 2d). In binary traits with heritability 0.1, population prevalence 0.1, a base cohort of with 120k effective samples, and a target cohort of 5,000 effective samples, the mean for the adjusted PRS in relation to case-only overlap is 1.61×10^−4^ (standard error = 2.11×10^−4^) (Fig 2b) and the mean for the adjusted PRS in relation to control-only overlap is -3.77×10^−5^ (standard error = 1.98×10^−4^) (Fig 2c). On the other hand, when unadjusted, the mean in relation to case only overlap is as high as 0.161 (standard error = 8.93×10^−4^), whereas the mean is 6.28×10^−3^ (standard error = 2.59×10^−4^) in relation to control-only overlap.

While EraSOR effectively eliminates inflation caused by inter-cohort overlap in all simulation scenarios tested in relation to heritable traits, false-positive results are still observed after EraSOR adjustment in non-heritable traits when there is a large degree of overlap (> 0.29). For non-heritable quantitative traits with base cohorts of 120k samples and target cohorts of 10k samples, if all target samples are also present in the base cohort, then we observe a false-positive rate of 87%, with a mean of 0.0028 (standard error = 1.14×10^−4^). This is likely caused by the fact that a key component of the mathematics underlying the EraSOR algorithm (described by Eq. 11 in Methods) includes an estimate of *h*^*2*^ in its denominator. Therfore, when the trait is non-heritable, Eq 9 may be unstable and lead to an error in the ErasOR adjustment. However, we recommend that polygenic risk score analyses should not be performed on traits with estimated *h*^*2*^ < 0.05 (see [1]) and, thus, in sufficiently powered applications of PRS, EraSOR should have strong performance.

One of the main assumptions of EraSOR is that there is no environmental stratification 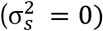. To investigate the robustness of EraSOR to model misspecification, we also performed simulations by incorporating UK Biobank samples with non-European ancestry and simulated different level of environmental stratification.

Overall, EraSOR is robust against model misspecification. For example, for quantitative traits with heritability of 0.1 and an environmental stratification of 0.3, the mean Δ*R*^2^ is -2.7×10^−3^ with standard error of 6.96×10^−4^ when the degree of overlap is 0.141 for target cohorts with 5000 samples and bas cohorts with 250k samples. EraSOR performs equally well for quantitative traits with different heritability, different level of environmental stratifications and cohorts with different sample size and overlap (see Supplementary Fig. 3-6).

### Robustness with diverse ancestry data

While the F_ST_ between the European and non-European samples (see Methods) is high (F_ST_ = 0.018), non-European samples accounts for only ∼5% of the UK Biobank population. Our results might therefore be dominated by the European samples. To understand how EraSOR performs in a more heterogenous dataset, additional simulations were performed using HapGen2 simulated genotype [22].

Using HapGen2 and the Finnish and Yoruban recombination maps from 1000 genome, we simulated 180k “Finnish” and 180k “Yoruban” samples. F_ST_ estimation from PLINK between the two population is only 0.00638, much lower than the reported F_ST_ > 0.1 African and European population [23]. This discrepancy is likely a result of the fact that the only difference in the simulations is the recombination map used (Supplementary Fig. 7) by HapGen2. Nonetheless, the HapGen2 simulation generates a dataset where the genetic signal is not dominated by one single population and allows us to better understand the performance of EraSOR in the presence of stratification.

Under the HapGen2 simulations, for quantitative traits with heritability 0.1, base cohort size of 250k, target cohort size of 5000, degree of overlap is 0.141 (all target samples also in the base cohort), environmental stratification 0.1, then the mean Δ*R*^2^ for the adjusted PRS is -1.01×10^−3^ (standard error = 9.41×10^−4^); the mean Δ*R*^2^ of the adjusted PRS is 0.0012 (standard error = 8.17×10^−4^) when the environmental stratification is 0.3, showing slight inflation. Nonetheless, when compared to the unadjusted PRS, which has mean Δ*R*^2^= 0.225, the inflation of the adjusted PRS is small.

One potential reason for the robustness of EraSOR may be due to the simplistic population structure of the simulated genotype. As we simulated the environmental stratification according to the population label, it is possible that by adjusting for PCs, the environmental stratification was fully adjusted for. While it is highly unlikely that environmental stratification is orthogonal to population genetic structure, we performed an additional simulation in which the population label was randomly assigned to the simulated genotype. This ensured the simulation of environmental stratification independent of population genetic structure and, thus, should not be capture by PCA adjustment (see Supplementary Methods and Supplementary Fig. 8).

Even when environmental stratification is simulated independently of the population genetic structure, EraSOR adjustments are still robust to different environmental structure. For quantitative traits with heritability of 0.1, base cohort size of 250k, target cohort size of 5,000 and environmental stratification of 0.3, the mean for the adjusted PRS is -6.08×10^−4^ (standard error = 9.68×10^−4^).

## Discussion

The recent advent of large-scale national and regional biobank projects, such as the UK Biobank [8], Japan Biobank [9] and FinnGen [11], have provided large resources of genotype-phenotype data ideal for conducting polygenic risk score analyses. However, this burgeoning generation of large data has led to an increased risk of inter-cohort sample overlap or relatedness, which can lead to inflated type 1 error. Due to privacy laws and practical concerns, it is usually impossible to identify overlapping samples or related samples across different cohorts. However, ideally researchers would be aware of the scale of the potential problem and have tools to mitigate against it. Therefore, here we reported on an investigation to evaluate the impact of inter-cohort sample overlap and relatedness in PRS analyses and developed a method to account for potential inter-cohort overlap and relatedness that does not require access to raw genotype data from the base GWAS.

We demonstrated that inter-cohort overlap results in a significant and often substantial inflation in the observed PRS-trait association, coefficient of determination (R^2^) and false-positive rate. This inflation can be high even when the absolute number of overlapping individuals is small if this makes up a notable fraction of the target samples. The inflation is noticeably more severe for binary traits with a small population prevalence when all the overlapping samples are cases. Therefore, PRS results will likely be misinterpreted unless inter-cohort sample overlap and close relatedness is properly accounted for.

Here, we developed the Erase Sample Overlap and Relatedness (EraSOR) method. EraSOR is designed to correct for inter-cohort sample overlap and relatedness using only summary statistics, without requiring any other information. The results of PRS analyses using EraSOR-adjusted GWAS results in the presence of sample overlap or relatedness was remarkably similar to those gained when the overlap was explicitly removed in most simulated conditions. EraSOR is also robust to mis-specification of the model, for example, when there is environmental stratification. While EraSOR does not fully adjust for the bias introduced by inter-cohort overlap for non-heritable traits when the degree of overlap is high, we recommend that researchers should not perform PRS analyses on non-heritable traits in any case [1]. EraSOR performs well for the majority of simulation scenarios tested here, which we believe reflect a large fraction of PRS studies.

Theoretically, as *ρ* from the bivariate LD score regression is assumed to be the phenotypic correlation [18], we can apply EraSOR in situations where the base and target cohorts measure different phenotypes. Based on LeBlanc’s equations [16], the spurious correlations caused by inter-cohort overlap and relatedness is a function of the phenotypic correlation. While this suggests that the impact of inter-cohort overlap and relatedness are likely to be smaller for cross-trait analyses, EraSOR adjustments may still be beneficial in these scenarios. Investigation of the performance of EraSOR in cross-trait analyses should be the subject of future work. Further research is also required to understand the performance and biases of EraSOR for applications in cross-trait studies and in its potential application to GWAS meta-analyses.

Our algorithm is not without any limitations. First, as EraSOR depends on the LD score intercept estimates for the adjustment, all assumptions of LD score regression also apply to EraSOR. For example, LD score regression assumes the level of genetic and environmental stratification is similar between the two cohorts [13], and if this assumption is violated, then it is likely that the bivariate LD score equation does not hold, which will lead to bias in EraSOR estimates. Moreover, due to reliance on LD score regression estimates, EraSOR only produces sufficiently accurate adjustments for application when both base and target cohorts have sample sizes greater than 1,000 and is only consistently accurate when both cohorts are greater than 5,000 samples. Nonetheless, despite its limitations, EraSOR is an ideal tool for application in settings in which there is known overlap in relation to large target samples and for sensitivity analyses in PRS studies. If the performance of PRS using the unadjusted and EraSOR adjusted summary statistics differs substantially, then this will act as a warning to the possible presence of inter-cohort overlap or close relatedness that should be adjusted for in order to obtain reliable PRS analysis results.

## Availability of supporting source code and requirements

|  |  |
| --- | --- |
| Project Name | EraSOR |
| Project Homepage | https://choishingwan.gitlab.io/EraSOR/ |
| Programming Language | Python (version 3.0+) |
| License | GNU General Public License version 3.0 (GPLv3) |
| Any restrictions to use by non-academics | None |
| Simulation Scripts | https://gitlab.com/choishingwan/sample_overlap_paper |

## Availability of supporting data and materials

All code used for this paper is available at **https://gitlab.com/choishingwan/sample_overlap_paper** and were implemented using nextflow (version 20.10.0 build 5430) [28].

## Abbreviations

PRS: polygenic risk score

## Additional files

Supplementary Table 1: Simulation results

## Authors’ contributions

Conceptualization, S.W.C. and P.F.O.; Methodology, S.W.C., T.S.H.M., C.H. and P.F.O.; Investigation, S.W.C.; Software, S.W.C.; Supervision, P.F.O.; Funding Acquisition, P.F.O.; Writing – Original Draft, S.W.C; Writing - Review and Edition, S.W.C., T.S.H.M., C.H. and P.F.O.;

## Acknowledgements

We thank the participants in the UK Biobank and the scientists involved in the construction of this resource. We thank Jonathan Coleman and Kylie Glanville for helpful discussions. This research has been conducted using the UK Biobank Resource under application 18177 (P.F.O.). This work was supported in part through the computational resources and staff expertise provided by Scientific Computing at the Icahn School of Medicine at Mount Sinai.

## Supplementary Materials

**Supplementary Figure 1.**
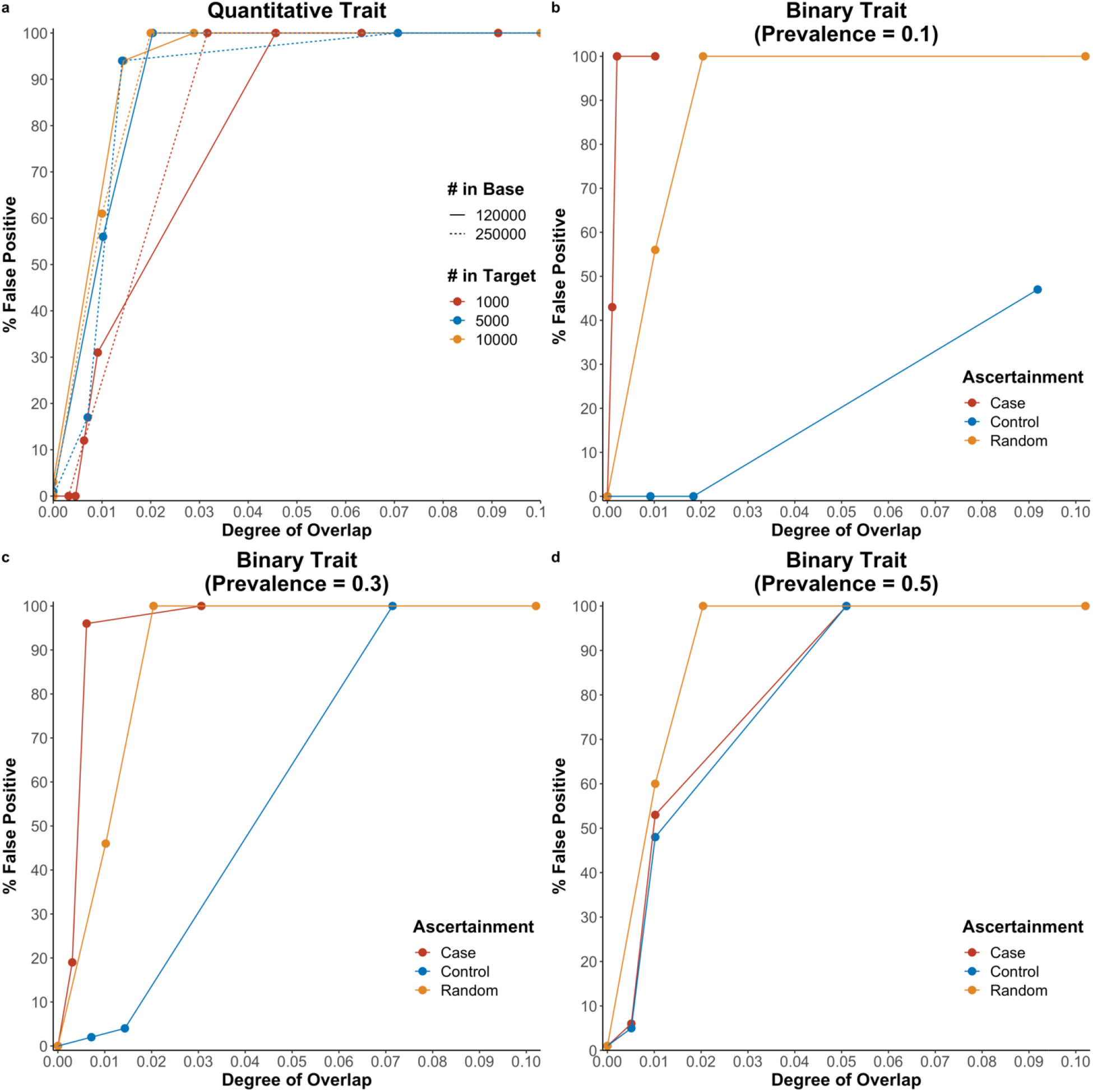
False positive rate corresponding to different level of sample overlap. Non-heritable phenotypes were simulated. X axis shows the degree of overlap, calculated as 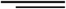 and the Y-axis shows the percentage of false positive (PRS P-value < 1×10-4). **a)** Quantitative traits with different cohort sizes **b)** Binary traits with population prevalence of 0.1 **c)** Binary traits with population prevalence of 0.3 **d)** Binary traits with population prevalence of 0.5.

**Supplementary Figure 2.**
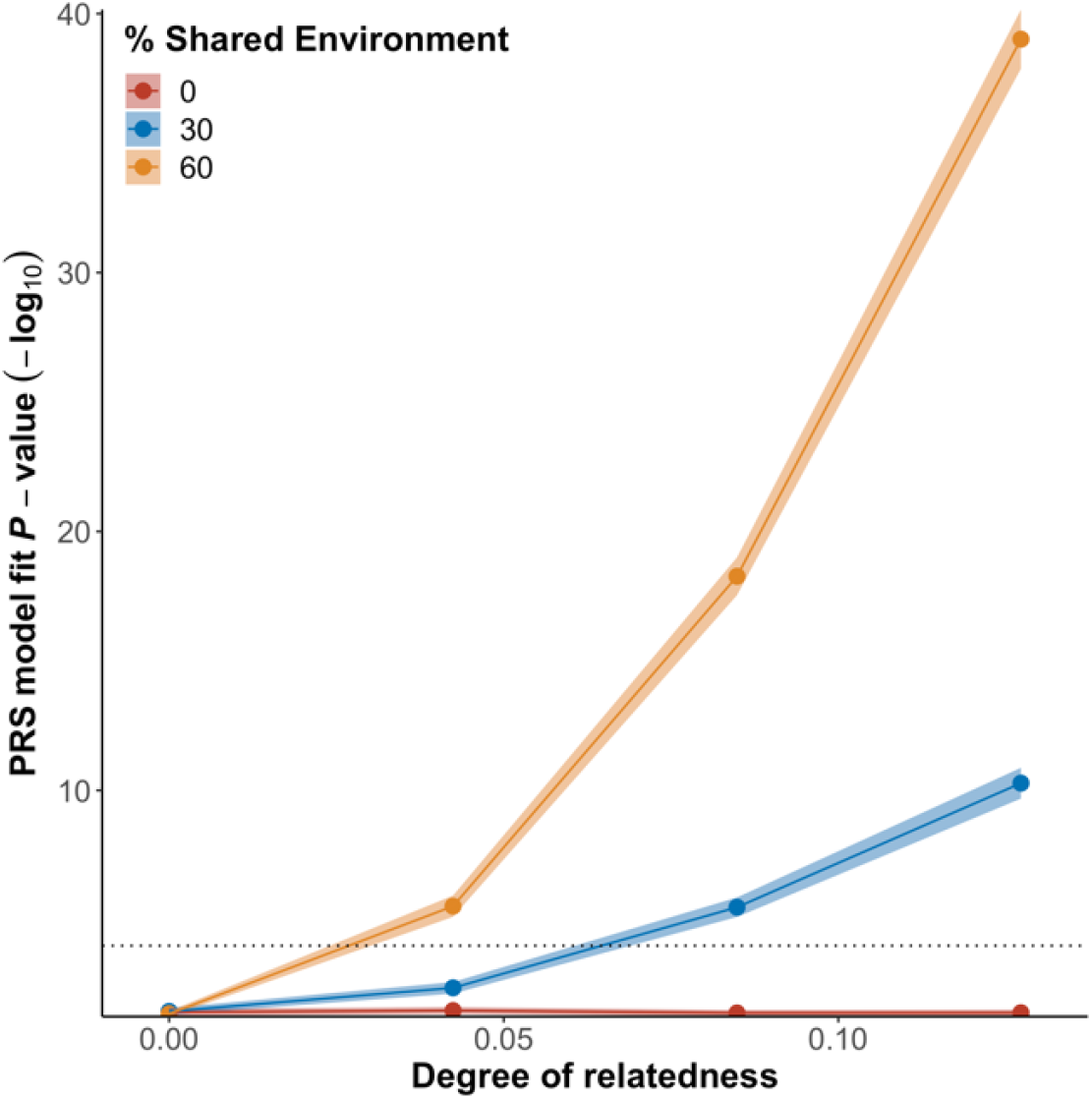
Effect of sample relatedness on performance of PRS for non-heritability phenotypes. The dotted line represents the significant threshold i.e., p-value < 1×10^−4^. Y-axis represents the -log_10_ transformed *p*-value of association between the PRS and the phenotype; the X-axis represents the degree of relatedness between the target cohort and the base cohort, calculated 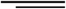 where is the number of samples in the target cohort that are first degree relatives to sample in the base cohorts. Shaded area represent the 95% confidence interval.

**Supplementary Figure 3.**
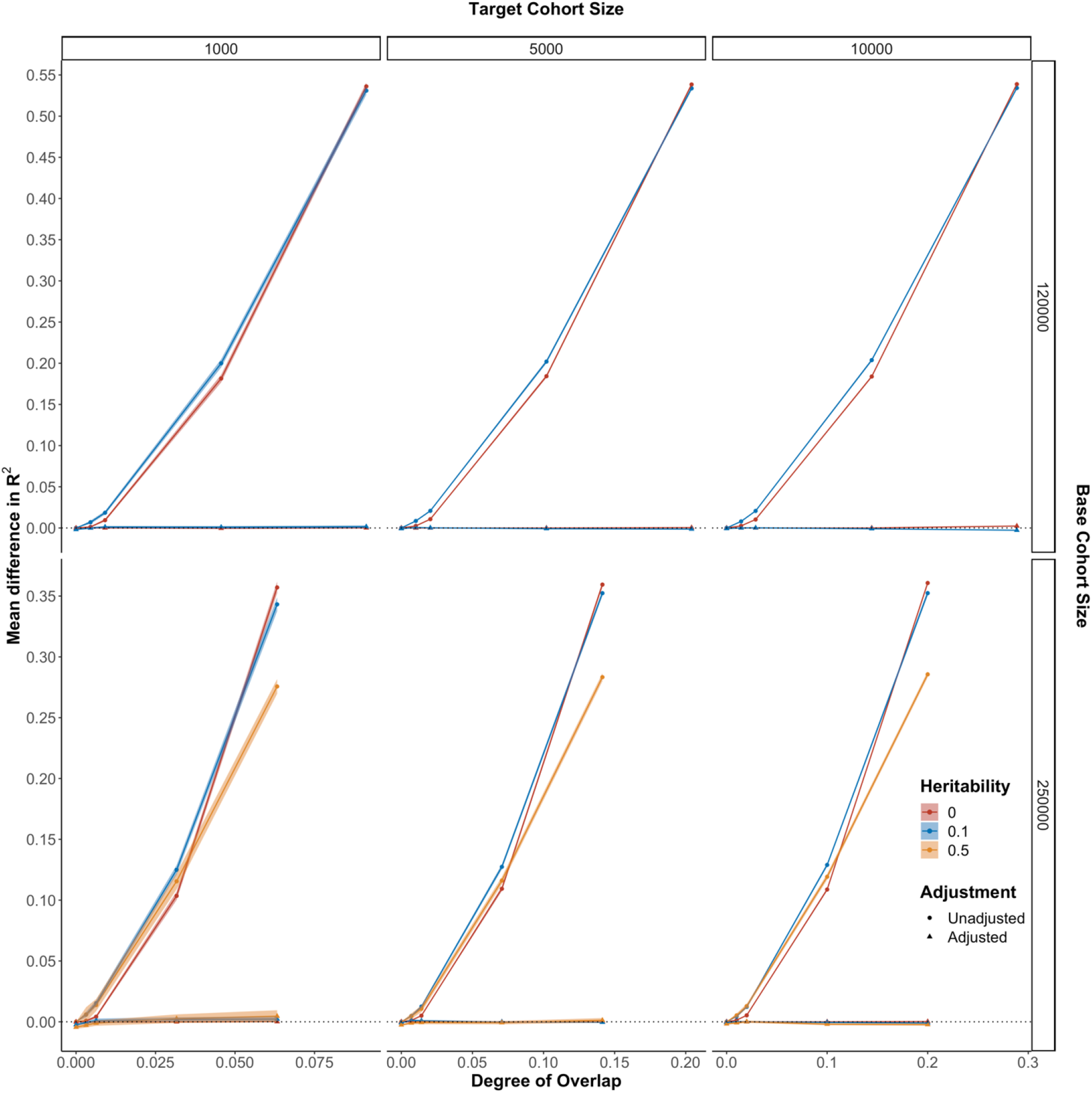
Comparing the performance of PRS using the EraSOR adjusted summary statistics and the unadjusted summary statistics for quantitative trait without population stratification. The X-axis shows the degree of overlap, and the Y-axis shows the mean difference between the observed R^2^ and the expected R^2^. Mean difference in R^2^ = 0 is represented by the black dotted line. Each row corresponds to different base cohort sizes, each column corresponds to different target cohort size and different colors correspond to different trait heritability. Performance of the adjusted PRS is indicated with triangle and performance of the unadjusted PRS is indicated with circle. Shaded area represents the 95% confidence interval, which tends to be small.

**Supplementary Figure 4.**
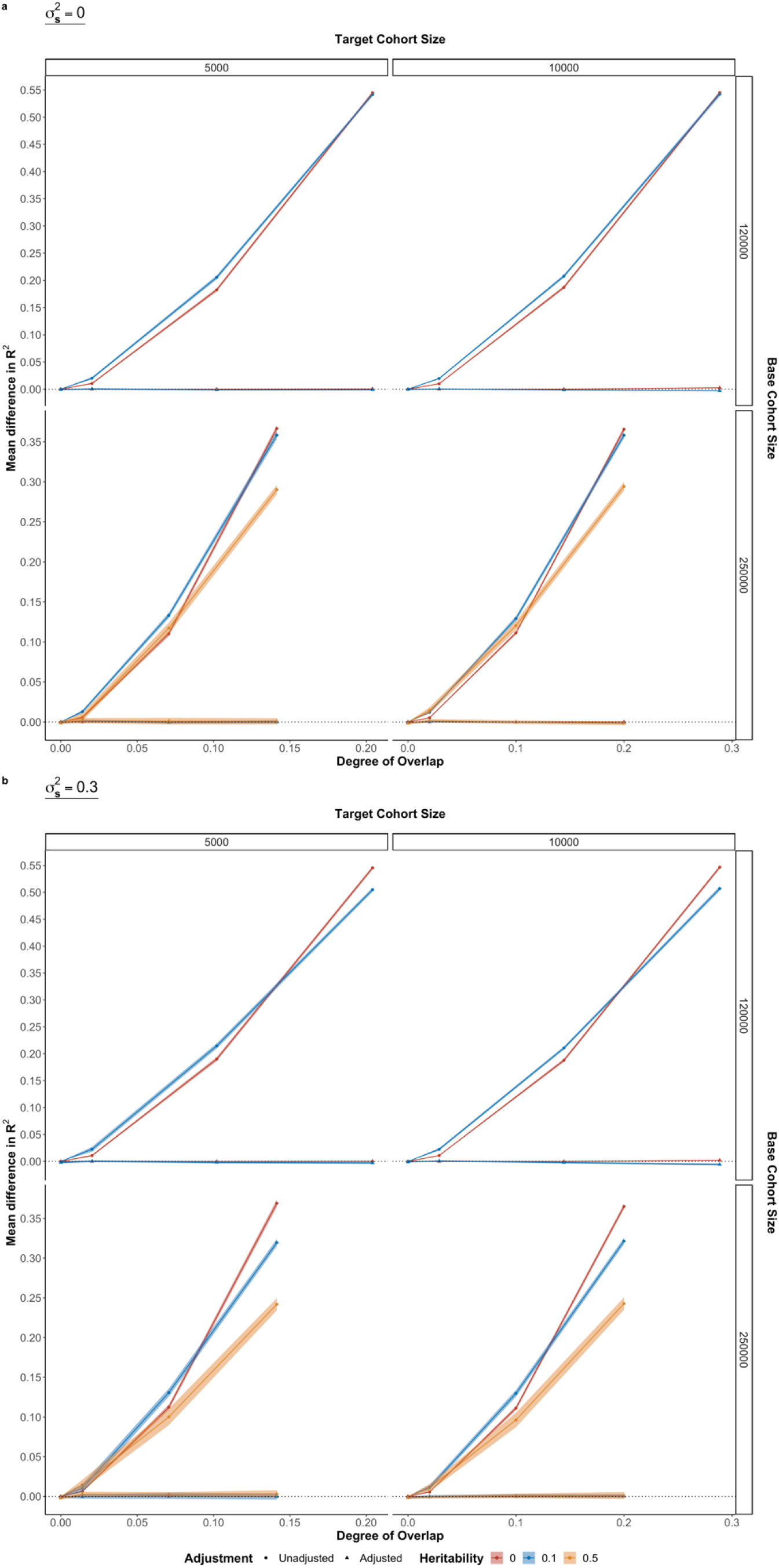
Comparing the performance of PRS using the EraSOR adjusted summary statistics and the unadjusted summary statistics for quantitative trait when there are population stratifications. Samples from non-European ancestries were included in this analysis. Different level of environmental stratifications was simulated: **a)** no environmental stratification **b)** environmental stratification = 0.3. Shaded area represents the 95% confidence interval, with different colours represent different simulated heritability. Results of the unadjusted PRS were represented with circle, while results of the adjusted PRS were represented with triangle.

**Supplementary Figure 5.**
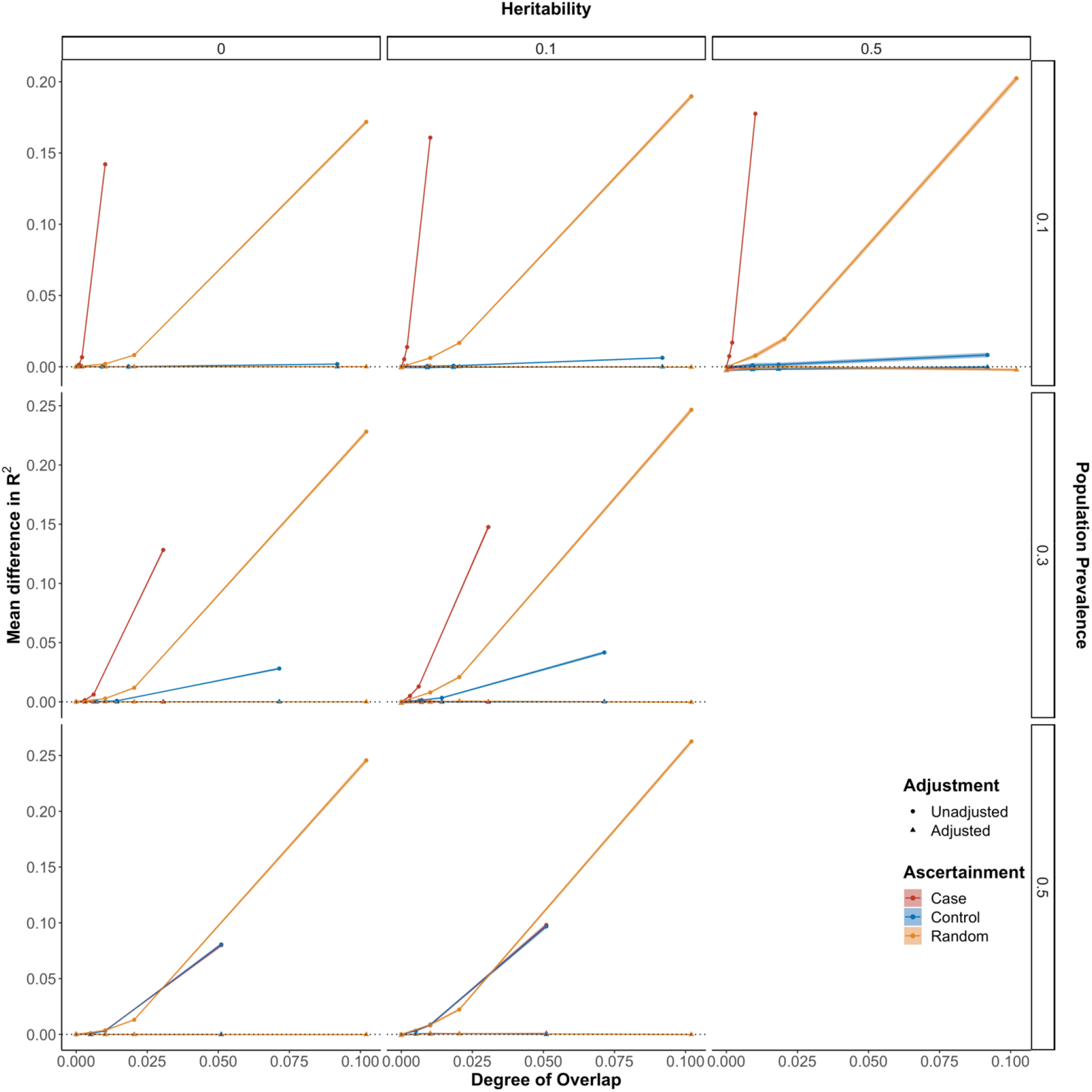
Comparing the performance of PRS using the EraSOR adjusted summary statistics and the unadjusted summary statistics for binary trait analyses. The X-axis shows the degree of overlap, and the Y-axis shows the mean difference between the observed R^2^ and the expected R^2^. Mean difference in R^2^ = 0 is represented by the black dotted line. Each row corresponds to different trait heritability, each column corresponds to different population prevalence and colors were used to represent different ascertainment of the overlapped samples. Performance of the adjusted PRS is indicated with triangle and performance of the unadjusted PRS is indicated with circle. Shaded area represents the 95% confidence interval, which tends to be small.

**Supplementary Figure 6.**
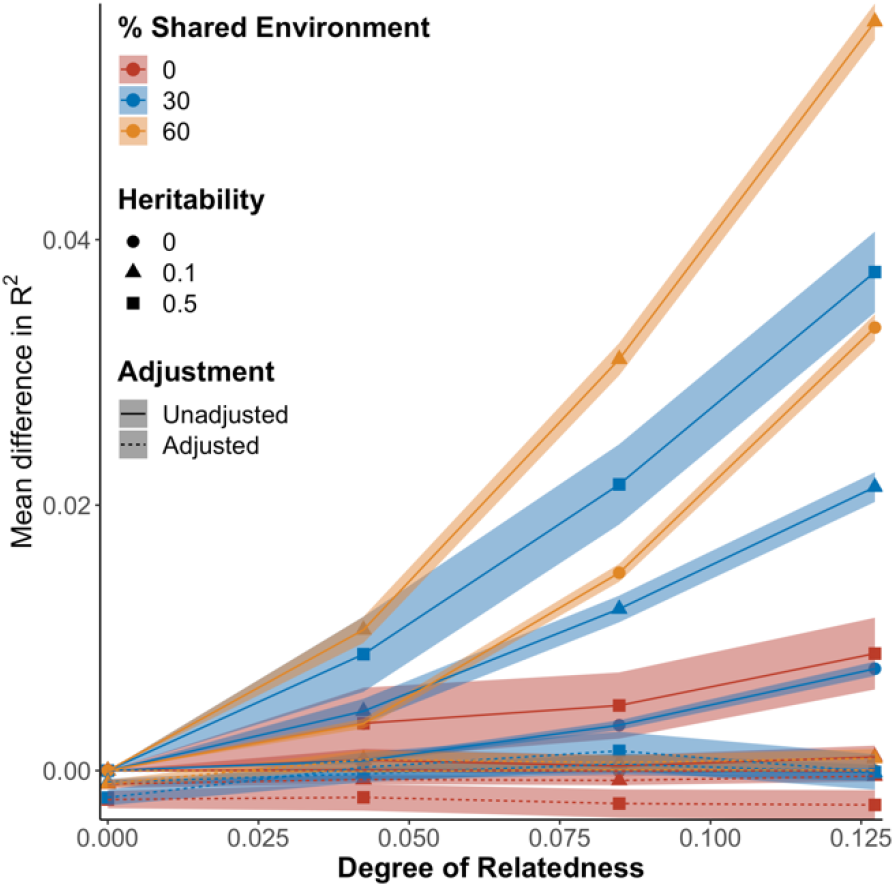
Performance of EraSOR when adjusting for related samples between the target cohort and the base cohort. The X-axis shows the degree of overlap, and the Y-axis shows the mean difference between the observed R^2^ and the expected R^2^. Shaded areas represent the 95% confidence interval. Different colours correspond to the amount of shared environmental contribution, and the shapes represents the heritability. Performance of EraSOR adjusted PRS are represented with the dotted line, and the performance of the unadjusted PRS are represented with the solid line.

**Supplementary Figure 7.**
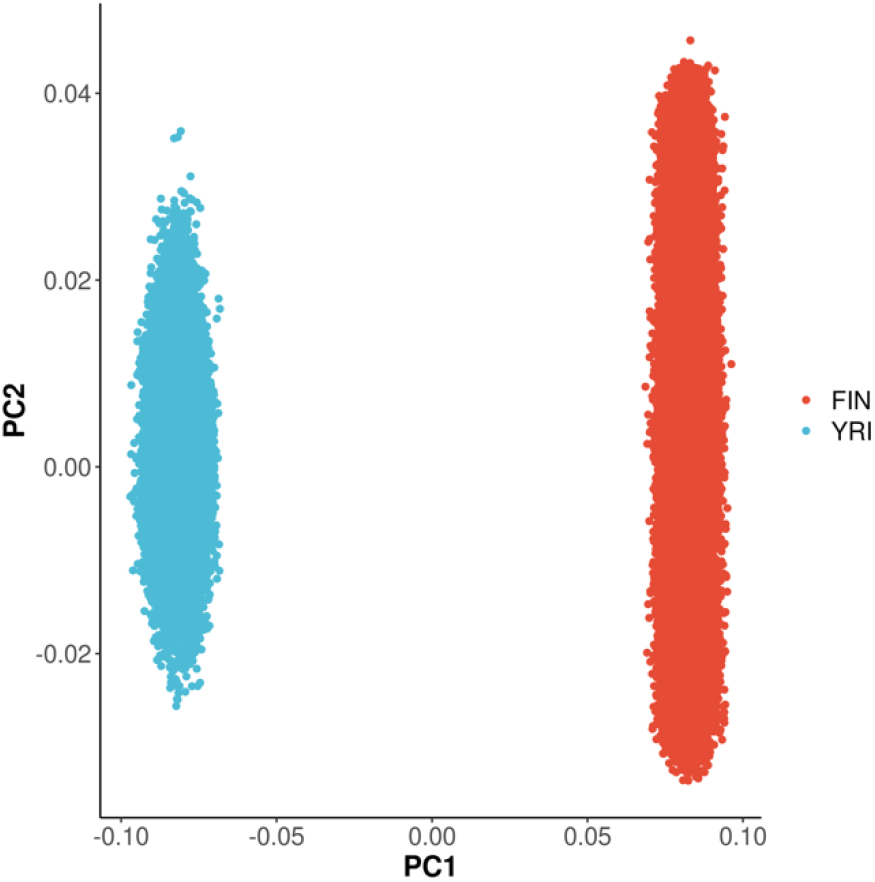
Principle Component plot for HapGen2 simulated data. Samples from the two populations were clearly separated by PC1, whereas PC2 does not contribute to the separation, suggesting that the simulated population structure might be much simpler than what would otherwise observe in real data.

**Supplementary Figure 8.**
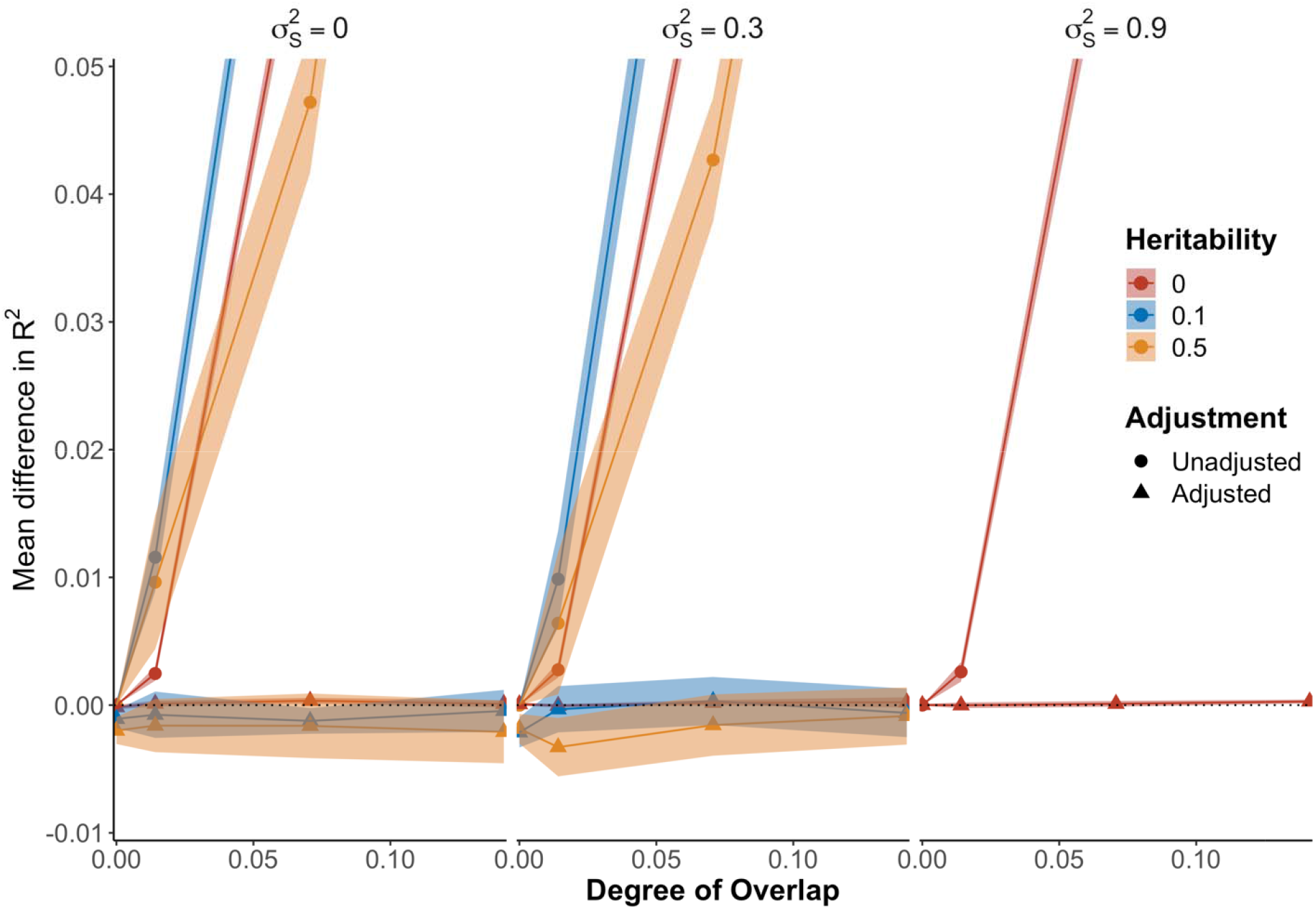
Performance of EraSOR adjusted PRS (triangle) against unadjusted PRS (circle) in HapGen2 simulations where phenotypes were stratified with agnostic environmental stratifications. The X-axis shows the degree of overlap, and the Y-axis shows the mean difference between the observed R^2^ and the expected R^2^. Shaded area shows the 95% confidence interval. Our result suggest EraSOR are robust against different environmental stratification () even when they are agnostic to the principal components.

## Supplementary Methods

### Simulate environmental stratifications agnostic to population structures

Similar to the main HapGen2 simulation, we used HapGen2 [22] to simulate 180k Yoruban and 180k Finnish samples using recombination maps from the 1000 Genomes Project [23]. 500 “Finnish” samples and 500 “Yoruban” samples were selected to calculate the LD scores using LDSC (v1.0.1) and flashPCA (v2.0) [24] was used to calculate the first 15 PCs of the data. 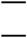 and 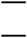 were randomly assigned to each individual, disregarding their simulated population.

The entire set of simulations were repeated 25 times.

